# Precon_all: A species-agnostic automated pipeline for non-human cortical surface reconstruction

**DOI:** 10.1101/2025.04.16.649072

**Authors:** R. Austin Benn, Ting Xu, Rogier B. Mars, Magdalena Boch, Lea Roumazeilles, Katja Heuer, Roberto Toro, Daniel S. Margulies, J.P Manzano-Patron, Paula Montesinos, Carlos Galan-Arriola, Gonzalo Lopez-Martin, Javier Sanchez-Gonzalez, Eugene P. Duff, Borja Ibañez

**Author notes:** Correspondence: R. Austin Benn.

## Abstract

Cortical surface reconstruction has transformed the study of brain morphology and cortical geometry in human neuroimaging. Here we present precon_all, an open-source structural imaging pipeline that generalizes reconstruction of the neocortical surface to multiple species. Adopting a modular framework that mirrors FreeSurfer’s recon-all in humans, precon_all produces surfaces for non-human species compatible with FreeSurfer and Connectome Workbench. Precon_all reduces the dependencies of surface reconstruction to an image with T1-like contrast and five user-defined masks. This removes the dependence on species-specific atlases and tissue priors, and in turn extends surface reconstruction to species where these resources do not yet exist. The pipeline supports a wide range of imaging conditions, including in-vivo and post-mortem data, and has thus far been used to reconstruct the cortical geometry of primates, carnivores, and ungulates across multiple sites, spanning both highly gyrified and lissencephalic brains. Precon_all can be automated for high-throughput processing, and its template construction module can build surface-based common spaces to enable group-level analysis of cortical morphometry in new species. In short, precon_all is a flexible, modular pipeline that enables the comparative neuroimaging community to study brain evolution through cortical geometry in an ever-growing number of species.

---

Understanding how brain structure and function enable behavior is a central goal of neuroscience. By studying the brain across species, comparative neuroscience observes this relationship at its natural extreme, where evolutionary pressures physically shape the brain, and thus the behaviors that define how a species interacts with its environment^1,2^. Historically, comparative neuroscience has relied on methods such as histology and gross dissection. While these methods have provided great insight into the organization of the brain across species, the time, labour, and expertise they require limit high-throughput sample collection and constrain the range of species that can be studied. Neuroimaging, by contrast, provides a relatively cost-effective, fast, and non-invasive method that enables rapid phenotyping and characterization of the whole brain^3^, making it ideal for broadly sampling the evolutionary space of brain organization^4,5^. Furthermore, the digital nature of neuroimaging lends itself to data sharing through initiatives such as the Primate Data Exchange (Prime-DE)^6^ and Digital Brain Bank^7^. These initiatives have the potential to bring us into an age of large-scale and high-throughput comparative neuroscience. However, for this to happen, the tools and analytic pipelines that have made large-scale neuroimaging studies of the human brain possible must first be translated to non-human species.

One of the early successes of human neuroimaging was the ability to recover the native geometry of the cerebral cortex through cortical surface mesh reconstruction^8–10^. Through these surfaces, the three-dimensional volume captured by magnetic resonance imaging (MRI) can be reconciled with the true biological structure of the cortex: a two-dimensional sheet folded in three-dimensional space. Cortical surface meshes are reconstructed as a set of triangles connected at shared vertices. The resulting surface provides a continuous representation of the cortical sheet that can be inflated, flattened, and resampled while preserving the cortex’s topology and spatial structure, providing greater flexibility for downstream analyses. This is especially valuable for studying aspects of cortical morphometry, such as cortical thickness and surface area, or topographically defined aspects of brain organization, such as gradients of connectivity^11,12^ or variations in receptor density^13^. Such versatility has made surface reconstruction an integral part of anatomical processing in large-scale human neuroimaging initiatives such as the Human Connectome Project^14^ and UK Biobank^15^.

Automated surface reconstruction in non-human species would mark a major step towards bringing comparative neuroimaging onto the same analytic footing as these large-scale human initiatives. Although this is already available in the rhesus macaque through pipelines like CIVET-Macaque^10^ and PREEMACS^16^, the broader value of surfaces in non-human research extends well beyond a single species. Surfaces have been used to relate cortical folding to evolutionary lineage^5^, to build sulcal^20^ and white matter atlases in newly studied species^17,18^, and provide the basis for novel geometric based cortical parcellations^19^. Beyond these descriptive applications, surfaces have further enabled quantitative cross-species approaches built on cortical surface models. These include surface-based registrations that quantify relative cortical expansion^20^ and areal relocation^21^, the direct assessment of functional homology^22^, and the construction of a cross-species connectivity space^23,24^. Common to all these studies is a fundamental requirement: a digital representation of the cortical surface for every species under study. However, current non-human surface reconstruction pipelines often require species-specific resources unavailable in species being scanned for the first time. To extend these advanced surface-based approaches to the ever-growing number of non-human species being scanned, a generalized, species-agnostic approach to surface reconstruction is needed.

Here, we present precon_all (https://github.com/neurabenn/precon_all), a cortical surface reconstruction pipeline specifically tailored for the non-human neuroimaging community. Designed to work with existing human neuroimaging software, precon_all requires only an image with T1-weighted-like contrast (that is, brighter white matter than grey, with dark cerebrospinal fluid) and five basic user-defined masks: a brain mask, left and right hemisphere masks, a non-cortical mask, and a subcortical mask. This straightforward setup allows precon_all to be adapted for use in most mammals, requiring minimal anatomical knowledge from the user. Its adaptability makes precon_all a well-suited tool for the initial characterization of cortical anatomy in species new to neuroimaging, and enables researchers to extend surface-based neuroimaging approaches to a wide array of species.

## Results

### General philosophy

Precon_all is an open-source pipeline that simplifies the process of surface reconstruction for non-human neuroimaging. Its aim is to generalize this process by minimizing dependence on species-specific anatomical priors such as cortical atlases and volumetric templates. Less-studied species and those being scanned for the first time often lack such priors, leaving researchers without a starting point to study their cortical geometry. Precon_all addresses this by adapting the human surface reconstruction pipeline FreeSurfer to forgo cortical labelling and run without a species-specific atlas or template. Its modular design preserves the quality control checkpoints and troubleshooting steps familiar to users of FreeSurfer, while outputs in both FreeSurfer and GIFTI formats ensure compatibility with the broader human neuroimaging ecosystem. To achieve this, precon_all relies on a small set of requirements that can be derived from the structural MRI image itself (Fig. 1).

**Fig. 1.**
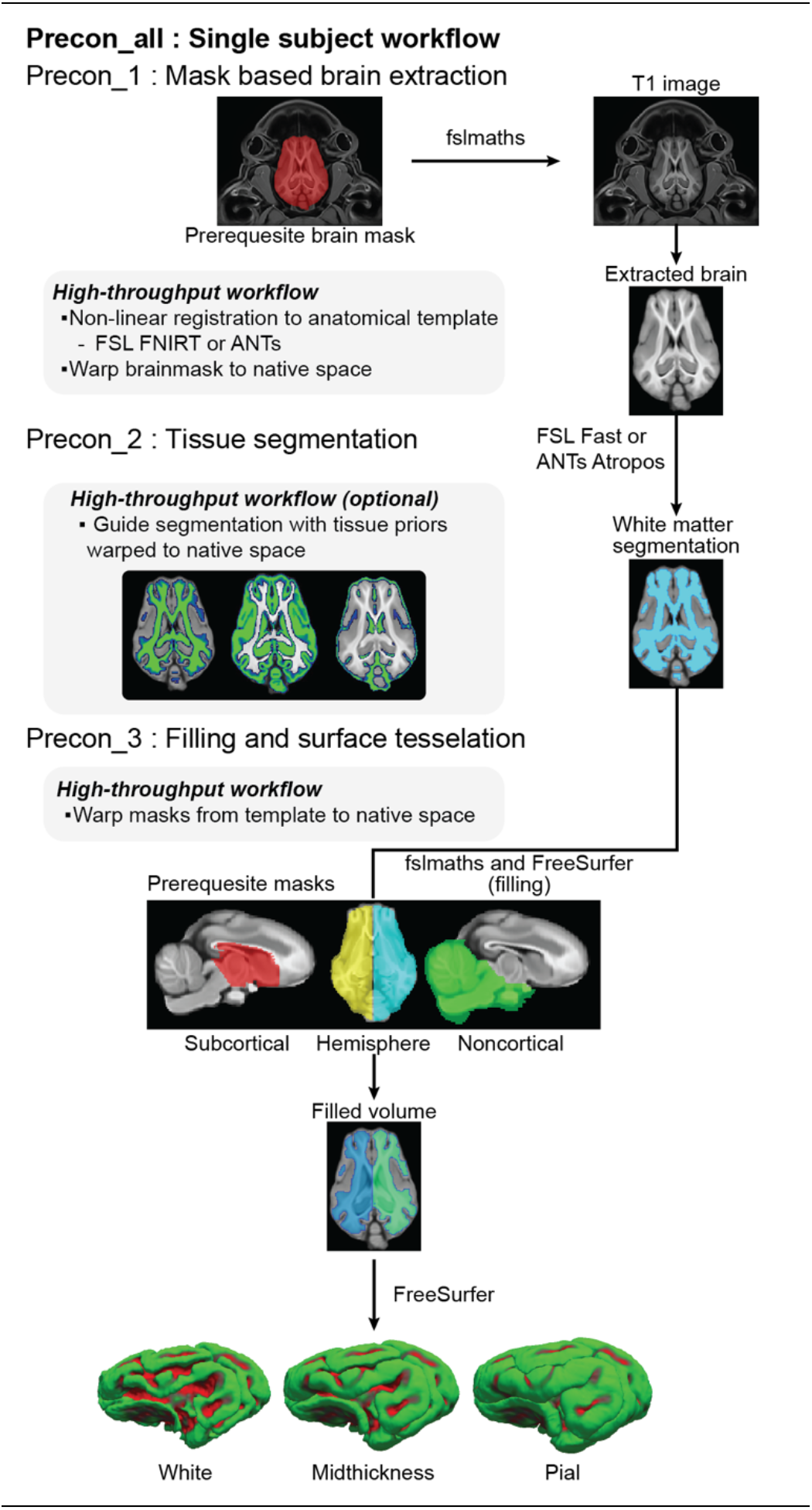
The precon_all pipeline. Precon_all’s modules: Precon_1 performs brain extraction, Precon_2 tissue segmentation, and Precon_3 filling and surface tessellation. Each module can be run individually as a breakpoint, or end-to-end by calling precon_all. Prerequisite masks and where they are used are shown for the domestic pig, and consist of the brain, hemisphere, subcortical and non-cortical masks.

First, precon_all requires a structural MRI image with T1-weighted-like contrast. This is because surface reconstruction in precon_all is performed with FreeSurfer, which first uses the highest intensity values in the image to reconstruct the white matter surface, and then expands outwards following the image intensity gradient to the boundary between the pia mater and cerebrospinal fluid. This contrast is not always available in non-human neuroimaging. Post-mortem samples, for example, are often acquired with T2-weighted or diffusion-weighted imaging. In such cases, we have found two approaches that produce suitable T1-like inputs. For T2-weighted images, we suggest a contrast inversion in which each voxel intensity is subtracted from the maximum within a brain mask and the absolute value is taken. For diffusion-weighted images processed with bedpostX, the root sum of squares of the f1 and f2 sample images produces an image with T1-like contrast (See utility script: *bedpostdir2pseudoT1.sh*). Throughout, we refer to actual T1 images as ‘T1-weighted’, inverted T2-images as “inverted T2” and those diffusion-weighted imaging derived T1s as ‘pseudo-T1’ images.

Second, precon_all requires the user to define five masks prior to surface reconstruction. These are a brain mask, left and right hemisphere masks, a non-cortical mask, and a subcortical mask (Fig. 1). The brain mask covers the cerebrum and is used in brain extraction. Hemisphere masks split the brain mask into its left and right components and ensure that FreeSurfer can reconstruct each hemisphere separately. An exclusion mask, the non-cortical mask contains regions such as the cerebellum and brainstem that are not part of the cortical surface. Finally, the subcortical mask fills the medial wall. Its posterior and inferior boundaries meet the non-cortical mask at the cerebellum and brainstem; it also fills the corpus callosum and the region beneath it, extending laterally to edge of the ventricles. Masks can be defined either in the native space of the structural image or, if available, in an anatomical template space. They can be created quickly using an open-source NIfTI viewer such as ITK-SNAP, whose interpolation tools allow each mask to be defined from a small number of slices. A detailed tutorial is provided through the following video tutorials (youtube.com/playlist?list=PLXhcGIatIV2U).

Once these requirements are met, precon_all’s command line interface can be accessed by running the *surfing_safari.sh* script. *Surfing_safari.sh* splits the surface reconstruction pipeline into three modules: brain extraction, segmentation, and cortical filling and surface tessellation (Fig. 1). Each module combines multiple open-source neuroimaging packages to prepare the file structure and pre-requisite images used by FreeSurfer to reconstruct the cortical surface (Fig. 1, Table S1). The output of precon_all mirrors the FreeSurfer directory structure: intermediate outputs of the segmentation and cortical filling can be found in the *seg* and *mri* directories, while the final unlabeled pial, white, and midthickness surfaces are contained within the *surf* directory. Finally, surfaces are output as FreeSurfer and GIFTI files enabling broader integration of non-human species into downstream workflows built on FreeSurfer, Connectome Workbench^25^, and Python packages like nibabel^26^.

### Adaptations for non-human data

Working with non-human neuroimaging data presents unique challenges due to the variable nature of the data. These variations can include acquisition parameters, different data orientations, anisotropic acquisition, field inhomogeneities, and the use of both in-vivo and post-mortem samples. Precon_all accommodates this variability. For example, in non-human primate studies it is common for species to be oriented in a “sphinx” position, where the animal is placed prone as opposed to supine in the scanner. Sphinx orientation can be corrected through FreeSurfer’s mri_convert. However, post-mortem studies vary more in their orientation, and manual reorientation tools such as Reorient (https://github.com/neuroanatomy/reorient) may be more appropriate. Precon_all accommodates orientation variability by reconstructing surfaces in the orientation of the input image. We would however, still recommend users use reorientation tools, before surface reconstruction to ensure that their images, and therefore the resulting surfaces, are aligned with standard neuroimaging orientation conventions (e.g., RAS, LAS, or LPS).

Other challenges, such as anisotropic acquisition or B0 field inhomogeneities, are handled directly by precon_all. In the case of anisotropic images, the data is resampled isotropically to the highest voxel dimension. Field inhomogeneities are corrected after denoising using the N4 algorithm as implemented in ANTs^27^. In our experience, the default N4 parameters generally work well. For data with stronger inhomogeneity fields, an additional bias field correction pass applied prior to running precon_all may improve performance as field inhomogeneity can significantly hinder segmentation performance. One option to overcome stronger inhomogeneity fields that may not respond to N4 bias field correction is the Analysis of Functional NeuroImages (AFNI) intensity normalization procedure, 3dUnifize. This can be run on the whole-head image prior to the pipeline, using a brain mask, via the utility script *IntensityNormalizeHeadImage.sh*.

Segmentation is its own challenge in non-human neuroimaging, as the contrast between grey and white matter tends to be lower, and tissue probability maps (priors) used to guide the segmentation of cerebrospinal fluid, white, and grey matter in human neuroimaging often do not exist. We, therefore, allow users to choose between two segmentation algorithms: ANTsAtropos^28^ or FSL’s FAST^29^, both of which can be run with or without tissue priors (Fig. 1, *precon_2*). Notably, both brain extraction and segmentation provide areas where users can adapt precon_all to the outputs of custom modules run externally. Precon_all can take a previously brain extracted image as input with the *-n* flag, and will accept alternate white matter segmentations (either hand drawn or externally derived) so long as it is saved in the *mri* directory as *wm_hand_edit.nii.gz.* These implementation choices enable precon_all to adapt to a wide array of data; for a summary of these software options see supplementary table S1, or the full precon_all development table (neurabenn.github.io/precon_all/precon_all_devtable.html).

Following segmentation, precon_all has the white matter mask required for FreeSurfer-based cortical surface reconstruction. As in the standard FreeSurfer workflow, this mask must first be prepared for tessellation. This involves separating the left and right hemispheres, removing the cerebellum and brainstem, and filling the subcortex and medial wall. FreeSurfer’s recon-all performs these steps via volumetric cortical and subcortical labelling based on a human atlas; a process which cannot be applied to species without an atlas. Precon_all addresses this by replacing FreeSurfer’s label-based operations with the hemisphere, non-cortical, and subcortical masks either manually defined for the individual being processed, or if later automation is desired in a species-specific template space (Fig. 1, precon_3). The hemisphere masks isolate each hemisphere; the non-cortical mask removes the cerebellum and brainstem; the subcortical mask fills the subcortex and medial wall. Together, these operations produce a combined white matter and subcortical volume, normalized to fixed intensities matching the inputs FreeSurfer expects, from which it reconstructs the white and pial surfaces natively. The resulting surface meshes are uniformly sampled as in line with human data (Fig. S1), and are saved in an output directory identical in structure to that of the standard FreeSurfer pipeline.

### Surface quality control and troubleshooting

Surface reconstruction with precon_all can occasionally fail. One of precon_all’s most common failures when working with a template space is inadequate brain extraction. To combat this, precon_all includes two template-based registration options for brain extraction. The legacy option uses FSL’s FNIRT to register an individual image to a template, after which the brain mask in the template space is warped to the individual to extract the brain. This method works best however when an initial *pre_extract.mat* linear FLIRT transformation matrix is provided, which a user can obtain through a rough registration of their anatomical image to the template space. Alternatively, brain extraction can be performed with the ANTs brain extraction workflow by passing the *-e* flag. This latter option can sometimes succeed where FNIRT based brain extraction has failed and can also be optimized through the same *pre_extract.mat* transformation used by the FNIRT workflow. Brain extraction tools are often highly species-specific, with several subfields having resorted to deep learning models to automate this process^30^. When brain extraction fails in precon_all, or a user has already run brain extraction through either a custom species-specific tool or manual segmentation, we recommend users premask their image to directly supply a brain-extracted image. Passing the *-n* flag to precon_all then allows this externally brain-extracted image to be used for the rest of the pipeline, bypassing the native brain extraction steps entirely.

Surface reconstruction errors after precon_3 are best detected through visual inspection, overlaying the white and pial surfaces on the anatomical image. This can be done in either Freeview, or automated quality control reports generated with the *precon_qc.sh* script. In particular users should look for incomplete expansion of the pial surface; the result of an inadequate white matter segmentation. As in human FreeSurfer runs, users should focus on areas such as the temporal pole and orbitofrontal cortex, though these areas may change based on the species being studied. These problems can be addressed using the same correction technique employed in FreeSurfer: manual refinement of the initial white matter mask and rerunning surface reconstruction (Fig. 2). In practice, this involves hand editing the white matter mask through a standard NIfTI viewer such as FSLeyes or

**Fig. 2.**
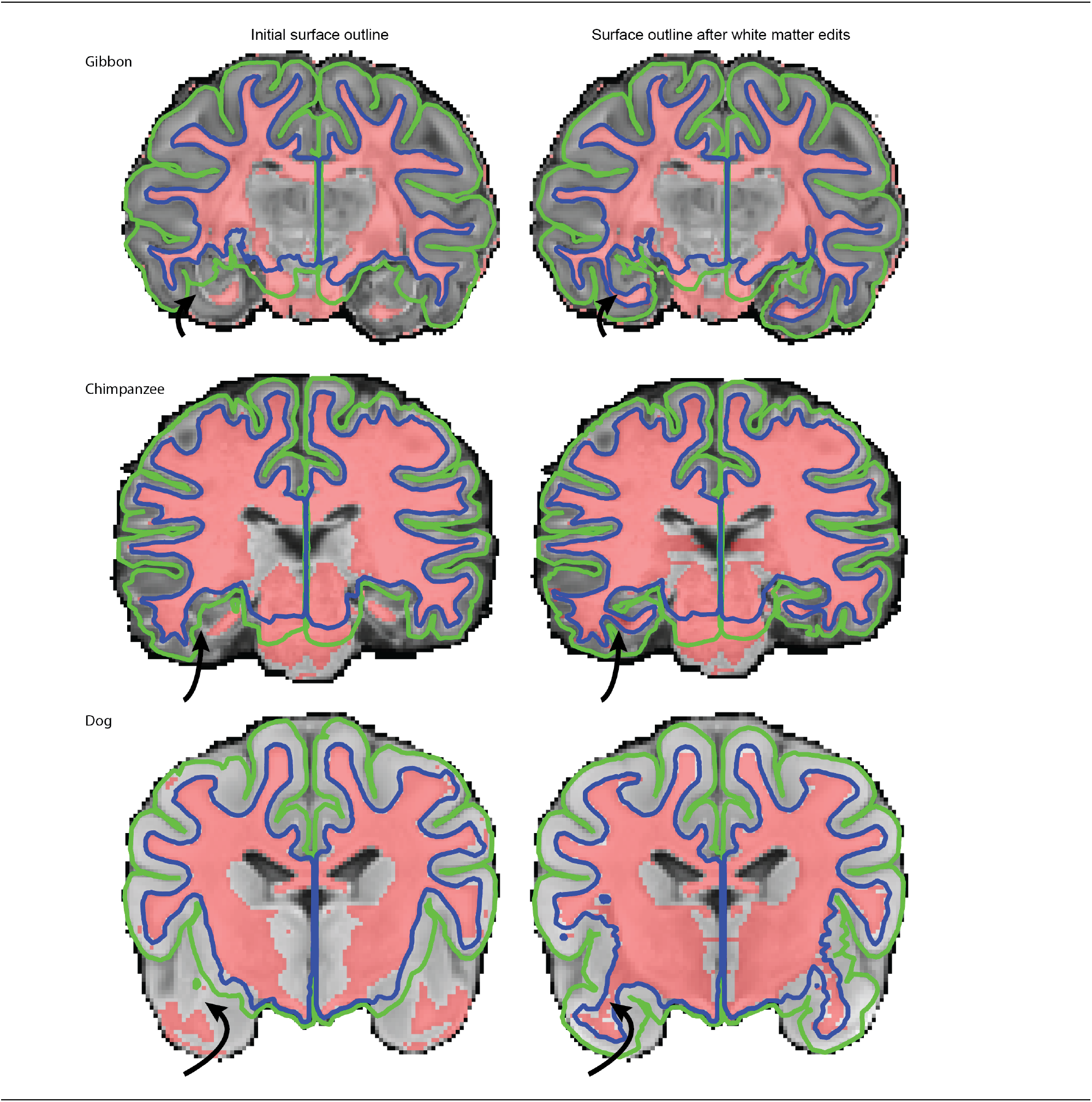
Surfaces are edited and updated as in FreeSurfer through manually editing the white matter mask. The gibbon image is a pseudo-T1 image derived from diffusion data to simulate contrast typical of a T1-weighted image (bright white matter, darker grey matter, dark CSF). This image is derived via the root sum of squares of the f1 and f2 bedpostX samples. The chimpanzee is a single T1 image from the National Chimpanzee Brain Resource, and the dog is a previously published T1 anatomical template image^53^. Arrows highlight areas where pial expansion was corrected following manual intervention through white matter mask correction.

ITK-SNAP, saving it in the *mri* directory as *wm_hand_edit.nii.gz* and rerunning *precon_3* (Fig. 1). This workflow also allows users to incorporate the results of alternative segmentation algorithms that may be better optimized for their species of choice, as any segmentation saved under this filename supersedes all others. This mirrors FreeSurfer’s standard quality control process, allowing researchers experienced with human pipelines to use the same workflow for non-human data. The only thing that changes is the species being segmented. As can be seen in Figure 2, final results are satisfactory.

In the presence of poor bias field correction and intensity normalization, the pial surface can over-expand, extending past the pia mater into the cerebrospinal fluid (Fig. 3). Precon_all includes bias field correction with the N4 algorithm^27^, though severe inhomogeneity may require additional processing. Whereas *IntensityNormalizeHeadImage.sh* is aimed at preprocessing the image before the pipeline is run, precon_all also ships with *intensity_afni_unifize.sh* for mid-workflow troubleshooting. This script normalizes the brain-extracted images from precon_1 with AFNI’s 3dUnifize command^31^, after which users can rerun precon_2 to improve pial placement.

**Fig. 3.**
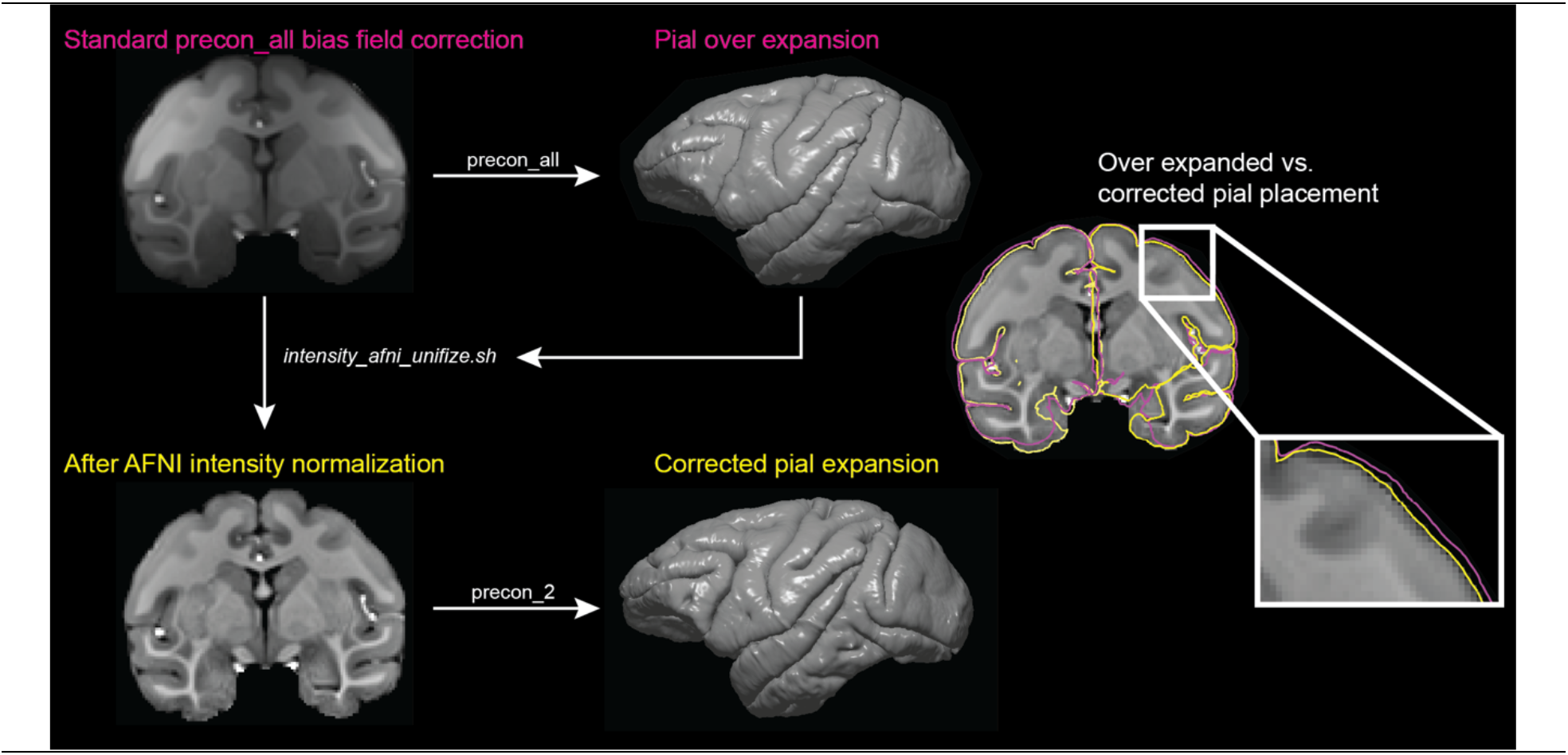
Insufficient bias field correction can cause the pial surface to expand past the pia mater into the cerebrospinal fluid. The workflow to fix this after an initial run of precon_all is to use the included utility script *intensity_afni_unifize.sh* followed by precon_2. **Top row (magenta):** an image where the default N4 bias field correction in precon_all was and the resulting over-expanded pial surface. **Bottom row (yellow):** the same image after intensity normalization, and the corrected pial surface generated by rerunning precon_2. **Right:** over-expanded (magenta) and corrected (yellow) placement of the pial surface.

### Demonstration across multiple mammalian orders and data types

The flexibility of precon_all positions it as a valuable tool for comparative neuroimaging, with demonstrated success across a wide variety of species (Fig. 4). It can process data from multiple branches of the phylogenetic tree, including primates, carnivores, and ungulates. One recent study used precon_all in 26 carnivores to describe sulcal morphology^17^, while the pig surface served as the foundation of a cross-species mapping between the pig and human^18^. Primates highlight precon_all’s versatility, reconstructing the cortical morphology of both highly folded brains of apes such as the gibbon^32^ and chimpanzee, to less gyrified species including lemurs, squirrel monkeys^33^ and galagos (Fig. 4).

**Fig. 4.**
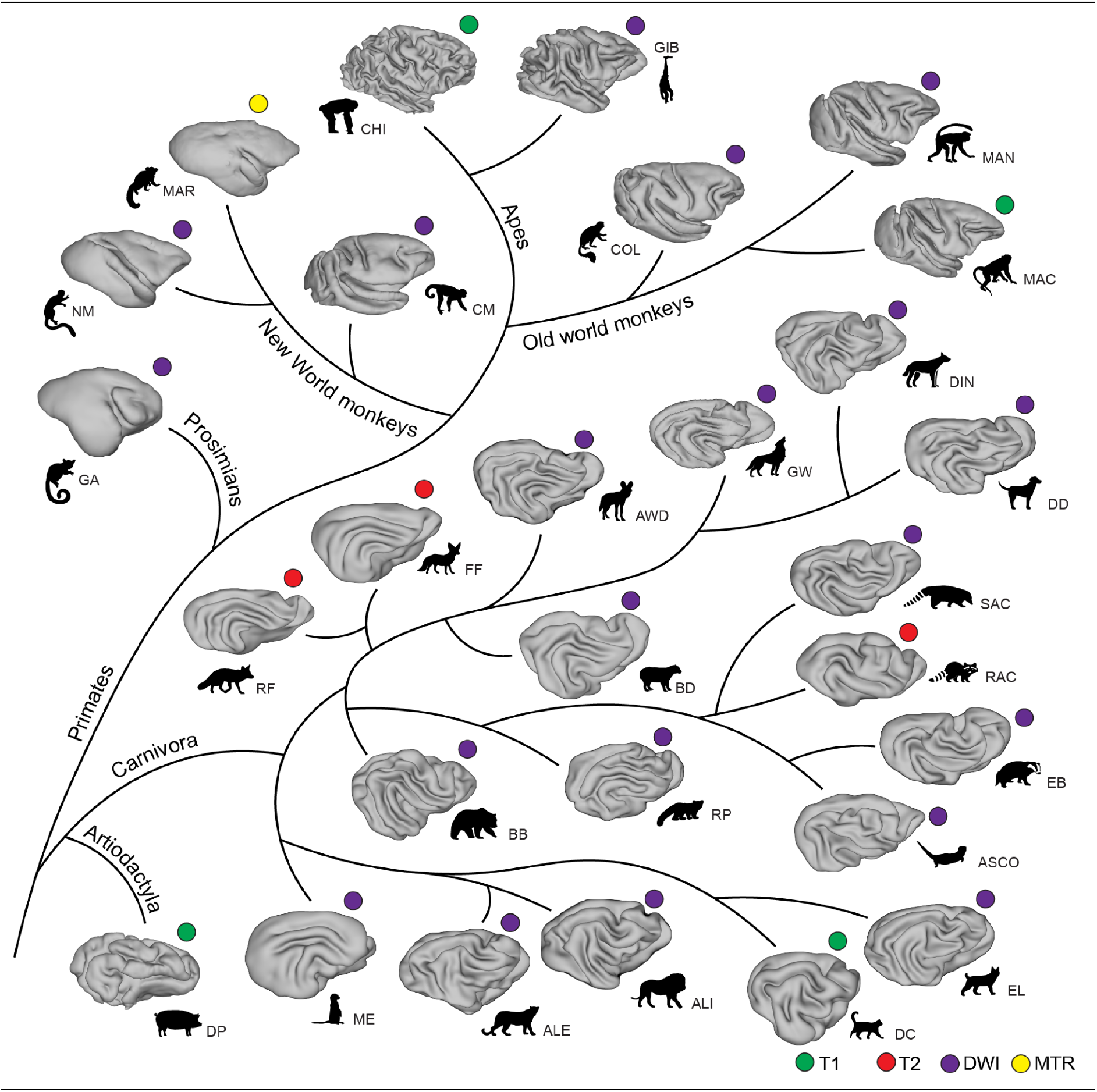
Cortical Surface Representations Across Species Representations of the right midthickness surface of 28 species reconstructed with precon_all. Acquisition modality is indicated by color, though as precon_all requires a T1 like contrast to run, only the T1 and proton density (PD) images were run on the native contrast. For T2 images, the contrast was inverted. For diffusion weighted imaging (DWI), pseudo-T1 images were generated by processing the data with bedpostX and taking the root sum of squares of the resulting f1 and f2 sample images. Abbreviations: ***Carnivora:*** ALI, Asiatic lion; ALE, Amur leopard; ASCO, Asian small-clawed otter; AWD, African wild dog; BB, brown bear; BD, bush dog; DC, domestic cat; DD, domestic dog; DIN, dingo; EB, Eurasian badger; EL, Eurasian lynx; FF, fennec fox; GW, grey wolf; ME, meerkat; RAC, raccoon; RF, red fox; RP, red panda; SAC, South American coati ***Artiodactyla:*** DP, domestic pig ***Prosimians:*** GA, Galagos ***New World monkeys:*** CM capuchin monkey; MAR, marmoset; NM, night monkey ***Old World monkeys:*** COL, colobus; MAC, macaque monkey; MAN, mangabey Great apes: CHI, chimpanzee; GIB, gibbon

Figure four also highlights precon_all’s ability to use real T1 and diffusion-weighted derived pseudo-T1 images derived from other modalities. In our example (Fig. 4), the domestic pig, domestic cat, and chimpanzee were processed using native T1 images, while the raccoon and red fox used inverted T2 images. The remaining surfaces were reconstructed using pseudo-T1 images derived from diffusion-weighted imaging. Furthermore, while not a T1 contrast, the marmoset surface was reconstructed based on a magnetic resonance transfer image, which at high resolution has a similar contrast to a T1-weighted image that was sufficient for precon_all to run. The surfaces we show here include a mix of in-vivo and post-mortem data acquired on different coils^34,35^, on a range of resolutions from 0.2 mm (marmoset) up to 0.8 mm (pig, chimpanzee).

The current version of *precon_all* is limited to a minimum voxel size of 0.2 mm, as pial expansion fails below this resolution. Future iterations will address this through a bespoke pial expansion algorithm that generalizes across small animal models scanned at high field. Because such an algorithm would lie outside FreeSurfer, it would require its own validation — a step the current pipeline avoids by inheriting FreeSurfer’s established human workflow. Despite this limitation, *precon_all* already accommodates much of the heterogeneity present in non-human neuroimaging and enables a broader set of species to bring surface-based analysis into their processing workflows.

### Scalability and group-level templates

As more non-human neuroimaging data becomes available through data-sharing initiatives like the Primate Data Exchange (Prime-DE)^6^, the Digital Brain Zoo^7^, Brain Catalogue (braincatalogue.org), the mammalian MRI dataset^36^, and the addition of non-human datasets to OpenNeuro^37^, there is a real need for high-throughput structural MRI processing. Precon_all addresses this by extending its single-subject workflow to multiple individuals. The prerequisite brain, hemisphere, subcortical, and non-cortical masks are first defined once in a volumetric template of the species being studied. Tissue priors, if available, can also be defined in this template space to guide segmentation (See Methods: Automating precon_all with anatomical templates). Each subject is then brain-extracted by non-linear registration to the template, with the same registration used to warp the prerequisite masks and any tissue priors into the subject’s native space. Quality control is supported by the *precon_qc.sh* module, which generates PDF reports covering brain extraction, surface outlines overlaid on the anatomical image, and a 3D rendering of the surface. Together with the template-based workflow, this allows individual surfaces to be reconstructed and validated across many subjects from a single set of template-defined inputs.

The template based workflow of precon_all has been previously used to reconstruct the individual surfaces of 50 pigs from a single site^18^. However, making the most of open non-human neuroimaging data requires scaling across sites, where scanners, acquisition parameters, and contrast differ. To test precon_all under different site conditions in the same species, we used data from the PRIME-DE^6^ data sharing initiative. Specifically, we reconstructed the surfaces of 29 macaques; nine from the Mount Sinai Philips dataset^38^, and 20 from the Oxford dataset^39^. Brain extraction run through ANTs in precon_1 was successful in 8/9 subjects from the Mount Sinai dataset, and 15/20 macaques for the Oxford dataset. Of the remaining six macaques where brain extraction failed, four were corrected by providing an affine matrix to initialize the registration. The remaining two macaques’ brain masks were edited by hand and passed to the utility script *manual_bet_fix.sh*, which uses the mask to extract the brain and updates the non-linear warps to template space. From there, users can call precon_2 via *surfing_safari.sh* to complete segmentation and surface reconstruction.

After recovering brain extraction in all 29 macaques, precon_2 was run to complete segmentation and surface reconstruction. In the Mount Sinai dataset, six out of nine macaque surfaces were reconstructed without the need for further intervention. The remaining three showed insufficient pial expansion in the medial temporal pole and parahippocampal gyrus, a known FreeSurfer issue typically omitted from surface quality control in humans^40,41^. Precon_all provides two ways to address this. The first, as in FreeSurfer, is manual editing of the white matter mask (Fig. 2). The second minimizes manual edits through region-specific thresholding of the white matter segmentation via the utility script *wm_regional_dual_thresh.sh.* This script lowers the segmentation threshold within a user-defined problem region (drawn in template space and projected into each subject’s space), allowing more white matter to be captured where needed. We applied this method to the three Mount Sinai macaques using a temporal pole mask defined in standard space, with the default global threshold and a lower threshold within the temporal pole. This recovered the surface reconstruction for all three macaques.

In the Oxford dataset, surface reconstruction initially failed in all 20 macaques, as the pial surface expanded beyond the cortex into the cerebrospinal fluid due to poor bias field correction. Using the *intensity_afni_unifize.sh* workflow (Fig. 3), we applied 3dUnifize to the brain extracted images in each precon_all directory and reran precon_2 with ANTs segmentation enabled. This successfully recovered cortical surface reconstruction in all 20 macaques from the Oxford dataset.

By design, precon_all reconstructs surfaces in the native space of each individual, supporting species that lack a volumetric template. Where a template does exist, however, group-level analysis becomes possible through spatial normalization to a common space. By preserving cortical topology, surface-based representations allow more accurate sulcal alignment than volumetric methods³⁴. Volumetric template creation has become relatively straightforward thanks to standardized pipelines in Advanced Normalization Tools^42^ and FSL’s MMORF^43^, but a surface equivalent for spatial normalization of non-human brains does not currently exist. Precon_all fills this gap through its template construction module, pet_sounds.sh (Fig. 5). To create a common surface space, the user runs precon_all on a volumetric template or single subject and supplies a list of quality-controlled precon_all output directories. Individual surfaces are then iteratively registered to the template space, after which the templates for registration are updated. This process occurs four times, after which a final registration template and average surface are generated. We demonstrate this with the 29 macaques from the Mount Sinai and Oxford datasets, constructing a macaque surface template that enables vertex-by-vertex comparison of cortical surface metrics across subjects.

**Fig. 5.**
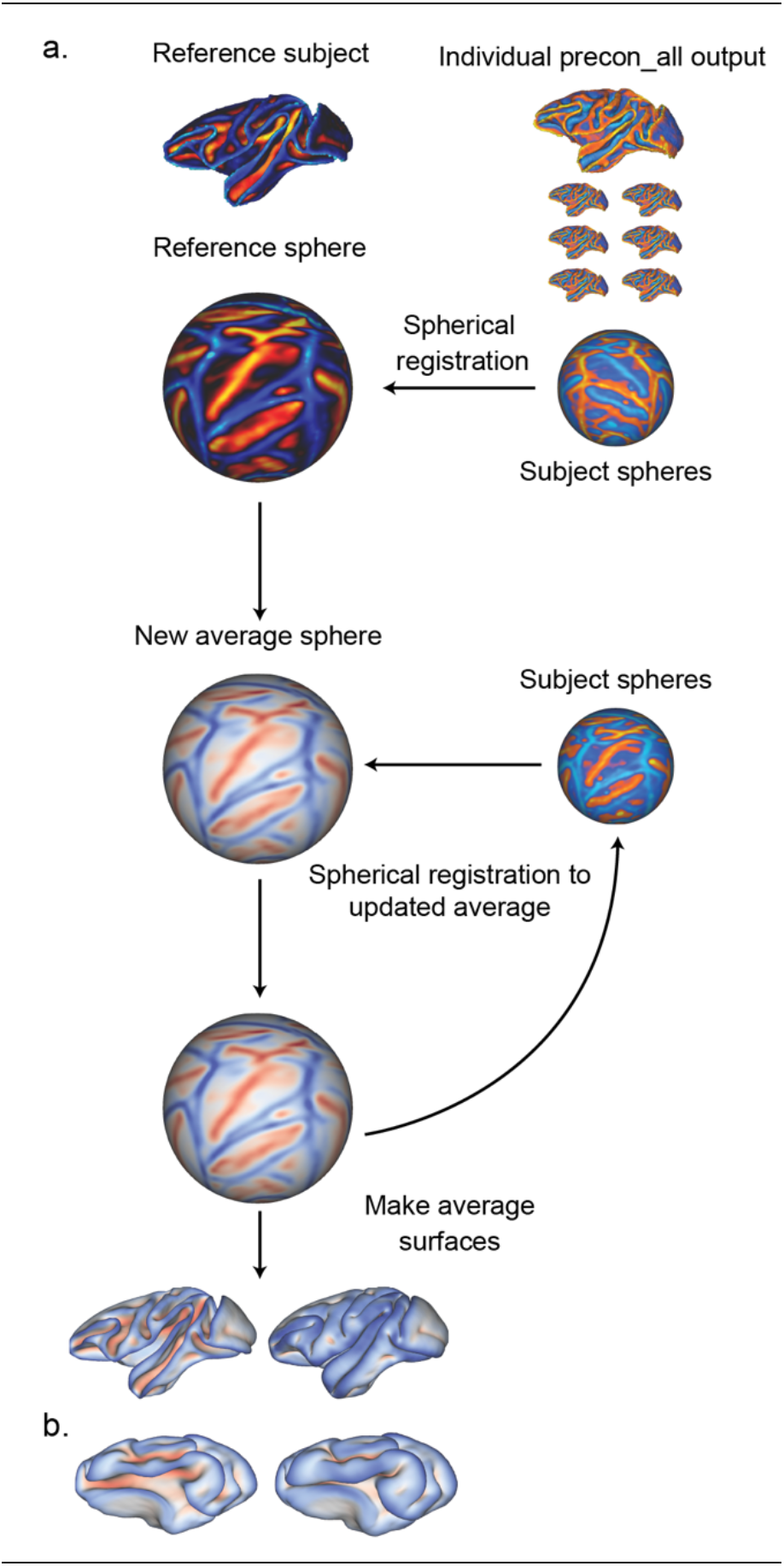
**a.** An overview of the process used to generate average surfaces for new species via the pet_sound.sh script of precon_all. This example represents the construction of an average surface using the NMTv1.2 as a reference space and 29 macaques sourced from the Mount Sinai Philips and Oxford datasets in Prime-DE^6^. **b.** An example group average surface using 50 domestic pigs^18^.

We then used this template space to assess how using a diffusion-weighted imaging derived pseudo-T1 as compared to a native T1 image could impact surface reconstruction (Fig. 6). To do this, we used the BigMac^44^ dataset which provides high resolution in-vivo T1 (0.5 mm), and post-mortem diffusion-weighted imaging (0.6 mm) acquired in the same animal. Individual surfaces were first derived using a standard precon_all run in the native space of each image. The surfaces of the real and pseudo-T1 images were then registered to the anatomical template space using the included utility script, *register_surfaces.sh*. This enabled a vertex-wise comparison of surface curvature, sulcal depth, and cortical thickness between in-vivo T1 and post-mortem pseudo-T1/DWI-derived surfaces. Curvature and sulcal depth showed high correspondence between surfaces (curvature: r = 0.87 left, 0.88 right; sulcal depth: r = 0.98 in both hemispheres), with minimal Bland-Altman bias for both metrics (≤0.2% of measurement range). Cortical thickness was moderately correlated between surfaces (r = 0.58 left, 0.60 right). Bland-Altman bias for thickness remained small (3.7% of range in the left hemisphere, 0.1% in the right), but limits of agreement were substantially wider than for curvature or sulcal depth (50 to 53% of range, versus 12 to 17%), indicating greater vertex-level imprecision in thickness estimates. We thus conclude that real and pseudo T1 images enable users to capture comparable representation of a species cortical geometry. However, studies aiming to assess cortical thickness in or across species should do so within a single imaging modality and refrain from comparing across them. More broadly, this example highlights how the native surfaces of precon_all can be spatially normalized, a critical step toward building surface-based multimodal pipelines like fMRIPrep³⁹ and HCP-like³⁸ workflows across a growing range of non-human species.

**Fig. 6.**
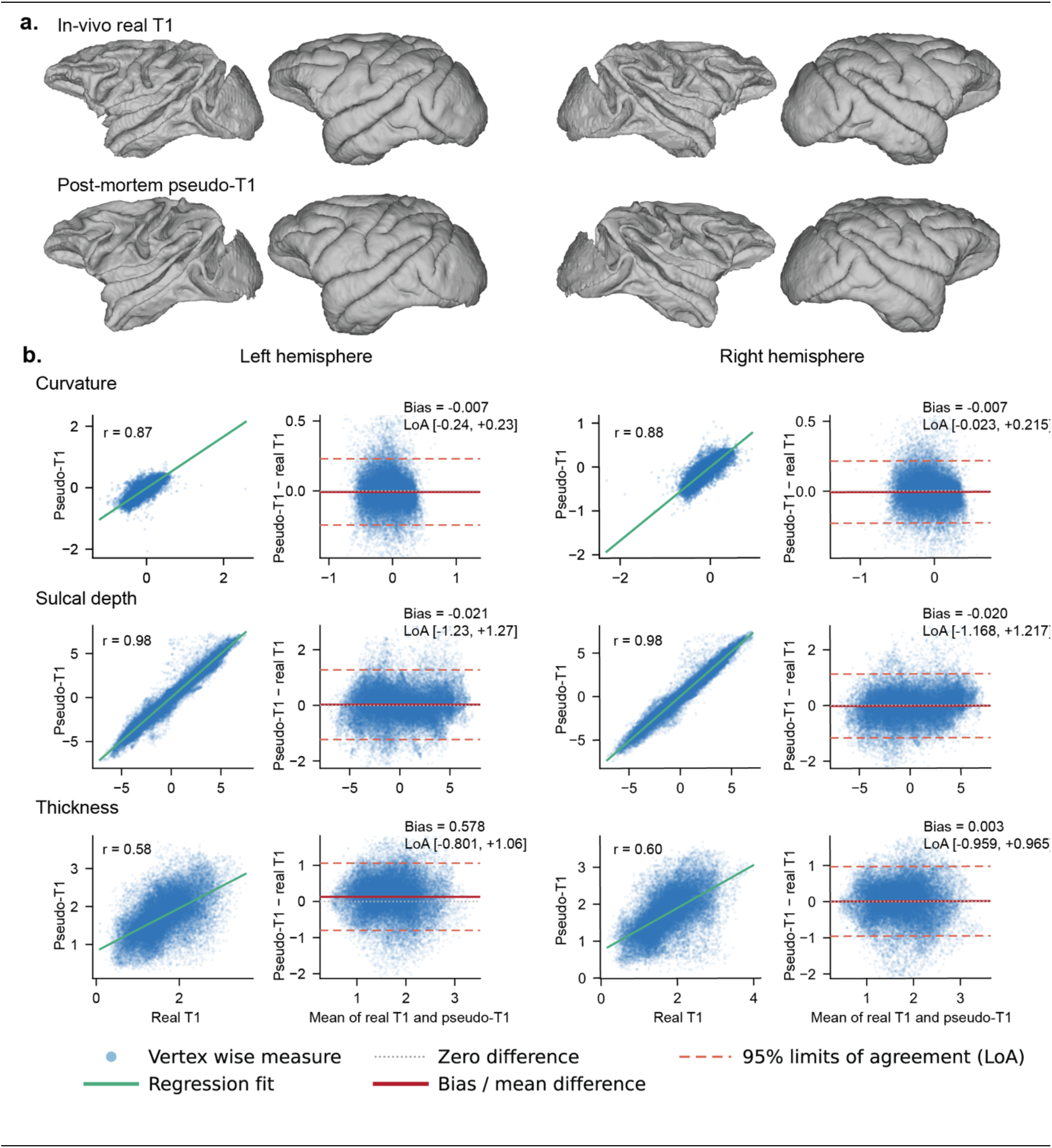
T1-weighted and diffusion-weighted-imaging-derived pseudo-T1 surface agreement from the same animal. **a.** Left and right surfaces from the BigMac^44^ dataset: in-vivo T1-weighted and post-mortem DWI-derived pseudo-T1 cortical reconstructions**. b.** Agreement between real and pseudo-T1 surface measures for the left and right hemispheres: vertex-wise correlation and Bland-Altman plots for curvature, sulcal depth, and cortical thickness. In correlation plots, the solid red line shows the linear regression fit. In Bland-Altman plots, the solid red line shows the mean bias and dashed red lines show the 95% limits of agreement.

### Implementation and availability

Consistent with our general philosophy, precon_all adapts established human neuroimaging software rather than introducing custom binaries. By relying on core functionality from each package, it remains compatible across a wide range of releases (see precon_all development table https://neurabenn.github.io/precon_all/precon_all_devtable.html). Precon_all was initially developed with a set of with minimum requirements that included FSL 6.0, FreeSurfer 6.0, ANTs 2.3.5, and Connectome Workbench 1.5.0. AFNI is an optional dependency, required only for the intensity-normalization utility script. However, our multisite scaling test in macaques (Fig. 5,6) ran using FreeSurfer 8.1.0, FSL 6.0.7.23, and ANTs 2.6.5, showing that precon_all can be maintained alongside the human neuroimaging software it is built on. Precon_all is open-source and available at https://github.com/neurabenn/precon_all. For users who prefer a self-contained environment, a Docker container with all required software pre-installed can be pulled from Docker Hub (https://hub.docker.com/r/neurabenn/precon_all). Alternatively, users can install the dependencies natively from their respective project pages (FSL: https://fsl.fmrib.ox.ac.uk/fsl/docs/; FreeSurfer: https://surfer.nmr.mgh.harvard.edu/; ANTs: https://github.com/ANTsX/ANTs; Connectome Workbench: https://www.humanconnectome.org/; AFNI: https://afni.nimh.nih.gov/), and then clone the precon_all repository and set a local precon_all path variable in their user profile.

Furthermore, the computing requirements of precon_all are no greater than those of its component human neuroimaging packages, making surface reconstruction feasible on modest hardware. Processing the 29 macaque scans from the Mount Sinai and Oxford datasets (0.5 mm anatomical resolution) on a single CPU core with 12 GB RAM on an AMD EPYC 7643 processor (2.3 GHz), mean total runtime for precon_all was 148.4 minutes (SD 26.6 min). Individual components of precon_all averaged as follows: ANTs brain extraction 19 min 34 s (SD 6 min 09 s); ANTs segmentation 12 min 30 s (SD 5 min 05 s); white matter filling 37 s (SD 16 s); and surface reconstruction 57 min 23 s (SD 14 min 44 s) for the left hemisphere and 58 min 21 s (SD 14 min 26 s) for the right. In sum, precon_all is openly distributed and runs with dependencies and hardware requirements accessible to most labs.

## Discussion

We have presented precon_all, a species-agnostic pipeline for cortical surface reconstruction. The pipeline builds on existing, proven standards in the neuroimaging fields and provides clear, robust adaptations to create compatibility with non-standard non-human data from a range of sizes, modalities, acquisition protocols, and qualities (Fig. 4). We have demonstrated the feasibility of precon_all and it has already been used in a number of published manuscripts^17,18,33,45,46^. Furthermore, the pipeline can run on local computers and HPC clusters, and includes utilities that enable scaling to high-throughput datasets. This pipeline opens up an important avenue for comparative neuroimaging to reach the analytic standards established in human work.

Data processing pipelines in neuroimaging are often highly modular, with specialized tools optimized for different steps such as image reconstruction, correction of acquisition biases, registration, subject-level statistics, and group-level inference. Most of these tools developed for human data require developer expertise and extensive modification to be used on data from non-human species^47^. However, some software tools are adapted or even optimized for processing non-human data. For example, end-to-end pipelines include the Rodent Automated Bold Improvement of Epi Sequences (RABIES) pipeline^48^, modified HCP pipelines for macaques^49^ and marmosets^50^, and the configurable pipeline for the analysis of connectomes (C-PAC) for macaques, and rodents^51^. However, it is also common for modules tackling a specific part of a processing pipeline to be adapted. For example, spatial normalization and volumetric template construction with ANTs has been done for the macaque^52^, pigs^18^, dogs^53^, and marmosets^34^, and mice^54^. Species-specific tractography protocols that plug in to FSL XTRACT^55^ have also been created developed for macaques^56,57^, and pigs^18^. While many of these adaptations suit the species they were built in, other packages like MrCat provide between-species statistical analyses that enable cross-species comparison^58,59^. It is important to identify the unique niche of precon_all in this ecosystem.

Precon_all takes minimally processed structural data and to reconstruct cortical surfaces. These surfaces can then be used for further processing. Such surfaces then support downstream analyses. For instance, cortical surfaces have been useful in integration of data across-modalities, such as transcriptomic and anatomical data^60^, macro-and microscale information^61^, and neurotransmitter data and computational properties of the cortex^13^. The 2D sheet is also helpful in identifying overarching principles of organization, such as connectivity gradients^11,12^, the influence of cortical geometry in cognition^62^, and the geometric constraints of functional activity patterns^63^. In comparative research, cortical surface representations have been used to guide cortical labelling in primates^32^ and carnivores^17^. They have also been used to investigate principles of cortical folding^5^, and investigation of principles of areal and connectivity changes across species^21^. Precon_all’s use in a number of these studies provides a strong proof of its application and diversity.

Comparative neuroimaging is going through a big data revolution. New datasets incorporating both more species, such as the Digital Brain Bank^7^, Brain Catalogue (braincatalogue.org), and the mammalian MRI dataset^36^ cover a broader range of species. At the same time, population-level data is being openly shared for individual species, including macaques through PRIME-DE^6^, rats through the StandardRat^64^, and mice through a multi-centre composite dataet^65^. With this comes the problem of scaling. As we have shown above, precon_all is both robust against very distinct brain geometries and scales well with increasing sample sizes from different datasets. Equally critical for such large studies is not just the reconstruction of cortical surfaces, but the spatial normalization of surfaces to a common template for cross-subject comparison and group-level inference. However, for most non-human species, no such surface-based template exists – one must be built for each species. Precon_all’s *pet_sounds.sh* template construction module fixes this, making the construction of surface-based templates a routine step in non-human neuroimaging. Thanks to recent data-sharing initiatives^6,7,37^, precon_all enables non-human surface based analysis to meet the scale of the data now available. Given this data availability, we hope precon_all empowers researchers to integrate surface-based analysis into their workflows at both the single-subject and group level. In this way, precon_all opens a path to build multimodal processing pipelines that facilitate cross-species translation, enabling like-for-like comparisons with their human equivalents.

The concept of generalization embodied by precon_all comes with trade-offs, as precon_all’s flexibility introduces several limitations. Although precon_all gives users multiple options to optimally reconstruct cortical surfaces (Fig. 1), the responsibility for quality control of the final surfaces rests with the individual user. Furthermore, precon_all does not perform cortical labelling; a feature standard in FreeSurfer, and many non-human primates (NHP) adapted pipelines^10,16^. However, this intentional design choice is also what permits precon_all to generalize surface reconstruction as it removes dependence on pre-existing parcellations that may not exist in less-studied species. From the surfaces output by precon_all, users can cortically label species being studied with comparative neuroimaging for the first time^17,18,45^(Fig. 4).

In sum, precon_all generalizes surface reconstruction through a flexible pipeline in a species-agnostic manner. Despite its flexibility, it also provides standardization, adopting gold-standard human neuroimaging software for use across the broader animal kingdom. We hope that the widespread adoption of *precon_all* will accelerate insights into how evolution shapes brain structure and function, with the cortical surfaces it generates serving as a foundation for future discoveries in translational neuroscience^17,18,45^.

## Methods

### Pipeline overview

Precon_all is built on FreeSurfer^9^, FMRIB Software Library (FSL)^66^, Connectome Workbench^25^, and Advanced Normalization Tools (ANTs)^67^. It provides a non-human neuroimaging pipeline capable of taking an image with T1-like contrast as input to produce white, pial, and midthickness cortical surface meshes. Precon_all recreates the outputs and directory structure of FreeSurfer’s recon-all^9^ so that users familiar with human neuroimaging can adapt their current human-based workflows to animal models with minimal effort. As with FreeSurfer’s recon-all, precon_all consists of a set of modules consisting of brain extraction, white matter segmentation, surface tessellation and topology correction. Unlike FreeSurfer, it does not perform cortical surface parcellation, only generating anatomical surfaces. This is a key distinction, as it enables precon_all to run in an anatomically agnostic environment, removing constraints of template-based brain extraction, and parcellation as part of the surface generation process. This decision was taken as cortical parcellations are often not available in lesser-studied species, or in the case when only a single MRI image is available for a given species. However, the surface outputs of precon_all do enable users to create their own cortical parcellations for species where cortical atlases are limited or missing^17,45^. The impact of precon_all is thus twofold as it provides the tools to study the cortical morphology of lesser-studied species, while also enabling diverse species to be resampled to a common dimensionality such that the surfaces derived from precon_all can drive horizontal translations across the branches of the phylogenetic tree^8^.

### Data sets and image acquisition

#### Pigs

Precon_all was run on the PNI50 template brain, a volumetric average of 50 *large white* domestic pigs^18^. All animals were scanned under anaesthesia (Ketamine 20 mg/kg, Midazolam 0.5 mg/kg, and Xylazine 0.2 mg/kg) in the prone position using a 32-channel cardiac coil. The T1 weighted 3D Flash Image was acquired at 0.8 mm isotropic resolution with the following parameters: TR 10 ms, TE 4.8 ms, Flip Angle (FA) 10°.

#### Marmoset

The anatomical surface was generated from the National Institute of Health Marmoset Brain Atlas^34^. To meet the resolution limits of precon_all, the original Magnetic Transfer Ratio (MTR) image, acquired at 0.15 mm isotropic resolution, was down sampled to 0.2 mm. Prerequisite masks were then manually drawn in the down sampled space, after which precon_all was run on the processed data.

#### Chimpanzee

A single T1-weighted image (0.8 mm isotropic) was taken from the National Chimpanzee Brain Resource (NCBR) of the subject Fritz. Prerequisite masks were drawn in ITK-Snap^68^, and precon_all was run with manual intervention occurring after the initial white matter segmentation.

#### Dog

A high resolution 0.5mm isotropic stereotaxic T1-weighted dog template was used to create the dog surfaces in Fig. 2^53^.

#### Data from the Digital Brain Zoo

The remaining primates (Mangabey, Colobus, Gibbon, Capuchin, Night Monkey, and Galago) and all carnivores were processed using data obtained from the Digital Brain Zoo^7^. This dataset encompassed 24 of the 28 species included in this study (Fig 4), consisting primarily of post-mortem diffusion-weighted images. Full acquisition details can be accessed through the digital brain bank ^7^ (https://open.win.ox.ac.uk/DigitalBrainBank/).

#### Macaque

The macaque surface template was created by first running precon_all on the National Institute of Mental Health Macaque template, version 1.2 (Fig. 5)^52^. Following this run, precon_all was run on 9 individual macaques obtained from the Philips Mount Sinai School of Medicine, and 20 Macaques from the University of Oxford. Data was accessed through the Primate Data Exchange initiative^6^. In both datasets, multiple T1 images were available for each subject. Surfaces were reconstructed using the average of each subject’s images, obtained by running the FreeSurfer command mri*_robust_template*.

#### The Philips Mount Sinai School of Medicine dataset

The Philips Mount Sinai School of Medicine dataset^38^ was acquired with the following parameters: Each subject was anesthetized with 1.2% isoflurane through the duration of the scan, and T1 weighted images were acquired in a 4-channel head coil at 0.5 mm isotropic resolution with the following parameters: TR 1500 ms, TE 6.93 ms, TI 100 ms, FA 8°.

#### The University of Oxford

The University of Oxford dataset^39^ was acquired as follows: Animals were anesthetized with isoflurane, on a 3T MRI scanner. T1-weighted images were acquired with a four coil with an MPRAGE sequence. Voxel resolution was 0.5 mm isotropic, TR 2500 ms, TE 4.01 ms, TI 1100 ms, FA 8°.

#### The BigMac dataset

Two images from the same animal were obtained from the BigMac dataset^44^: an in-vivo T1-weighted image from the associated PRIME-DE folder, and a post-mortem diffusion-weighted image.

In-vivo T1 image: The T1 image was acquired with an MPRAGE sequence with 0.5 mm isotropic resolution, TR 2.5 s, TE 4.01 ms.

Post-mortem diffusion data: Diffusion data was acquired with 128 gradient directions, at 0.6 mm isotropic resolution, TR 10 s, TE 24.5 ms.

For full details regarding the BigMac dataset see its associated documentation: https://pages.fmrib.ox.ac.uk/amyh/bigmacdocumentation/

### Macaque surface template construction

Template construction was performed with the *pet_sounds.sh* module. First, the reference space for template construction was defined by running precon_all on the National Institute of Mental Health Macaque Template (NMT), version 1.2^52^. Next, all 29 individual precon_all runs from the Mount Sinai and Oxford datasets were spherically registered to the NMT surface space using the FreeSurfer command *mris_register*. The aligned surfaces were then averaged to create an updated surface registration template, and this process was repeated over four iterations in total (Fig. 5), after which a final registration template and average surface were generated.

### Generating T1-Like Contrast for Non-T1 Images

Precon_all requires a T1-like contrast for surface reconstruction, though many of the examples presented here (Fig. 4) were not acquired using native T1-weighted images. Specifically, T1-weighted images were only available for species such as the pig, macaque, domestic cat, and chimpanzee, which allowed these datasets to be processed directly without additional steps. The Magnetic Resonance Transfer (MTR) images from the marmoset provided a sufficiently similar contrast to T1-weighted data, enabling precon_all to run without further modification.

For datasets acquired with T2-weighted or diffusion-weighted images, a T1-like contrast was created using two different approaches. For T2-weighted images, we inverted the contrast by multiplying all voxel intensity values by -1, and then added the absolute value of the new minimum to ensure that the highest original values became the lowest. This transformation was followed by applying a brain mask to set cerebrospinal fluid (CSF) and background voxels to zero.

For diffusion-weighted images, the T1-like contrast was derived through standard preprocessing using FSL’s FDT and BedpostX pipelines. This process is shipped with precon_all through the utility script *bedpostdir2pseudoT1.sh*. Briefly, this script creates a pseudo-T1 image by calculating the root sum of squares of the *samplesf1* and *samplesf2* output by BedpostX. Both methods resulted in contrast sufficiently similar to T1-weighted images for precon_all to run successfully (Fig. 4)

### Assessment of mesh uniformity

Mesh uniformity of precon_all outputs was assessed in nine macaques from the Mount Sinai Philips dataset accessed through the PRIME-DE and nine subjects from the Human Connectome Project (HCP). Surfaces were reconstructed with precon_all for the macaque dataset and with FreeSurfer and the HCP structural pipeline for the human dataset. For each subject and each surface, we calculated the vertex areas of the midthickness surface using the HCP *Connectome Workbench command -surface-vertex-areas*. The gifti file was then used to calculate the total surface of each mesh, which was divided by the number of faces expected to give a perfectly uniform mesh. Vertex areas were then back-projected to their corresponding faces, and the mean absolute error between each face’s actual area and the ideal uniform-mesh area was calculated as a measure of deviation from uniformity.

### Comparison of T1-weighted and pseudo-T1 derived surfaces

To assess the reliability of surface reconstruction across imaging modalities, we used the BigMac dataset^44^, which provides in-vivo T1-weighted imaging (0.5 mm isotropic) and post-mortem diffusion-weighted imaging (0.6 mm isotropic) acquired in the same animal. A pseudo-T1 image was derived from the post-mortem diffusion data by computing the root sum of squares of the f1 and f2 bedpostX samples (see utility script: *bedpostdir2pseudoT1.sh*). Cortical surfaces were independently reconstructed for both the in-vivo T1 and the post-mortem pseudo-T1 using standard precon_all runs. The resulting surfaces were then registered to the group-average surface template constructed with the *pet_sounds.sh* script based on the 29 Mount Sinai and Oxford macaques using the *register_surfaces.s*h utility script. Curvature, sulcal depth, and cortical thickness were resampled into this common surface space, enabling vertex-wise comparison between modalities. For each metric, we computed the Pearson correlation coefficient between the T1-and pseudo-T1-derived maps for each hemisphere, and performed Bland-Altman analysis to assess systematic bias and limits of agreement.

### User Guide

Our aim here is to describe how users can use precon_all to create cortical surface meshes of their animal model. With examples from across the animal kingdom including non-human primates, ungulates (Pigs), and Carnivores, we describe the pre-requisite masks needed to run precon_all in each species, the outputs generated, and how they can be down sampled to establish a common dimensionality for horizontal translation across species. While precon_all can be run in a single subject, we also show how precon_all can be automated and used to create population-based average surfaces (Fig. S2), enabling spherical registration, and resampling of data across subjects in a common surface space specific to each species.

#### Prerequisite masks

Precon_all depends on the definition of 5 user-defined masks, including a brain mask, left and right hemisphere masks, a non-cortical mask (cerebellum and brain stem), and a subcortical (medial wall) mask (Fig. 1). Each of these masks can be hand drawn, and defined in either the native space of the volumetric image, or defined in a volumetric template space permitting the automation of cortical surface meshes. For a full tutorial on defining these asks see: youtube.com/playlist?list=PLXhcGIatIV2U. For a single subject run, allow us to use an example of a domestic pig. We first ensure the data has a T1-like contrast, and then define the prerequisite masks in a folder called “masks” in the current working directory of the volumetric image:

#### The brain mask

The brain mask (*brain_mask.nii.gz*) consists of a binary brain mask containing the entirety of the cerebrum. To make this mask users should define a mask which covers the entirety of the cerebrum.

#### The hemisphere masks

To comply with FreeSurfer, the left and right hemispheres of the brain are treated separately by precon_all. Accordingly, the brain mask is split into its left (*left_hem.nii.gz*) and right (*right_hem.nii.gz*) components so that each hemisphere is masked separately. This mask can be made by splitting the brain mask through the *fslroi* command. Users should ensure that the final masks retain the header geometry of the original brain mask; if needed, a linear registration with nearest-neighbour interpolation via FSL’s flirt can be used to place the images back into the original space.

#### The non-cortical mask

The non-cortical mask (*non_cort.nii.gz*) consists of the brainstem and cerebellum and is used to remove structures prior to the tessellation of the 3D surface. This mask can be treated as a general exclusion mask and anything covered by it will be removed from the image prior to surface tessellation.

#### The subcortical mask

To define the subcortical mask, we suggest drawing a mask that includes the corpus callosum at its superior border, and posteriorly and inferiorly extends to the non-cortical mask. Along the lateral axis the border is defined by the boundary of the temporal white matter and the ventricular atrium, and the anterior boundary in these lateral borders extends to include the striatum and pallidum. This subcortical mask *(sub_cort.nii.gz)* serves to fill in the subcortex and also demarcates the medial wall in the final surface. These masks can be defined using open source nifti-image viewer such as ITK-snap or BrainBox (https://brainbox.pasteur.fr) and mask definition can be sped up through automated interpolation methods.

#### Automating precon_all with anatomical templates

Precon_all can be fully automated for new species if the prerequisite masks are defined in the space of an anatomical template. Recently, anatomical templates for a wide array of species have been generated, in part thanks to automated processing via the template construction scripts provided in Advanced Normalization Tools^42^. To automate precon_all for a new species, a new folder with the name of the species is created in the standard subdirectory of the precon_all home directory. Within this directory, there is a copy of the brain-extracted template and two subdirectories named “*extraction”*, and *“fill”* and the optional *“seg_priors”* directory. Here we provide an example file structure using the domestic pig (Fig. S2)^8^.

#### The extraction directory

The extraction directory contains the files associated with brain extraction (Fig. S2). Within it, there should be a copy of the whole-head and brain-extracted anatomical templates as well as the brain mask.

#### The fill directory

The fill directory contains the hemispheric masks, as well as the non_cortical, and subcortical masks (Fig. S2).

#### The seg_priors directory

An optional directory that allows users to provide segmentation priors for the segmentation steps called on later by precon_all (Fig. S2). Having defined the prerequisite masks in the space of an anatomical template, their placement in the specified directory structure permits precon_all to be fully automated from brain extraction to surface tessellation on multiple subjects (Fig. S2).

#### Running precon_all

Precon_all’s modular workflow provides quality control breakpoints that allow for each part of the surface generation pipeline to be run independently to attain optimal surfaces. The command line interface of precon_all is familiar to those who have worked with human neuroimaging data and FreeSurfer’s recon-all, as the following modules can be called: precon_all, precon_1, precon_2, and precon_3^9^. As in FreeSurfer’s recon-all, calling precon_all runs the full pipeline, including brain extraction, segmentation, filling, and tessellation of the cortical surface. Running sub-modules 1–3 of precon_all runs separate steps independently such that manual edits can be made to optimize the output of the final surfaces. The following paragraphs describe the components of each precon_all module:

#### Precon_1

Running precon_1 only extracts the brain. In the context of a single subject, precon_1 consists of simply masking the image with the user-defined brain mask. However, the power of precon_1 is that when used with an anatomical template space it automates brain extraction across species. This can be done in one of two ways with non-linear registration of the individual anatomical image to the template space, followed by the warping of the template brain mask to the individual subject space. The default option of precon_all uses FSL FNIRT^69^ for this operation, however users passing the *-e* flag to their command will instead use ANTs brain extraction and SyN registration instead. It’s reasonable to run precon_1 separately from the rest of the pipeline to troubleshoot this first step, as it can be assessed if a population-specific template may be needed to optimize registration to an anatomical template space. Furthermore, precon_1 might be used in lower-resolution scans where surface generation may not be possible, but spatial normalization is required. Users with a T1-weighted image that is already brain extracted, for example via U-net extraction^30^ can simply add the “-n” flag in the command line and run precon_all -n, as this will set up the appropriate directory structure and then invoke precon_2 such that the rest of the pipeline runs seamlessly.

#### Precon_2

Precon_2 segments the extracted brain image setting up the files needed for cortical “filling” and surface tessellation in precon_3. The key output of precon_2 is the white matter segmentation of the brain-extracted T1 weighted image. Users can choose to segment the image using either FSL’s FAST^29^ or ANTs Atropos^28^.

#### Precon_3

Surface generation in precon_all is based on FreeSurfer; specifically the command, “*mris_make_surfaces”*^9^. Precon_3 sets the stage for “*mris_make_surfaces”* by filling the subcortex of the white matter segmentation and removing the areas specified in the “non_cort.nii.gz” mask.

In the filling process, the T1-weighted image is first intensity normalized, and the binarized white matter segmentation is used to mask out the white matter in the T1-weighted image. The binary white matter mask is multiplied by 110 and then placed back into the T1-weighted image, as FreeSurfer expects the white matter to have a fixed value of 110 to demarcate the upper bounds of the image intensity gradient used to place the white and pial surfaces. Similarly, the area contained in the *“sub_cort.nii.gz”* masks is removed and replaced with a fixed value of 250 in line with FreeSurfer’s expectations. Once these intensity value substitutions have been made, surface prep is begun, removing the areas circumscribed by the *“non_cort.nii.gz”* mask, such that we are left with only the cerebrum. The left and right hemispheres are separated as a final step before tessellation using the *“left_hem.nii.gz* and *right_hem.nii.gz”* after which *“mris_make_surfaces”* is then run to generate the white and pial surfaces using FreeSurfer. Finally, the midthickness surface is generated via expansion of the white surface to 50% of the cortical thickness measured via the cortical ribbon, and all outputs are converted with *mris_convert* using the *“--to-scanner”* flag which produces GIFTI files compatible with *“connectome workbench”*^19^.

### Optional preprocessing

Precon_all is robust to a large range of data quality and variance in acquisition parameters (Fig. 4). However, as with all processing pipelines, it can be limited by the quality of the input it receives. While precon_all includes many built-in features, there may be times when additional preprocessing by the end user can help the pipeline complete successfully. This may include brain extraction with an external algorithm; for example, brain extraction in macaques is likely better performed using deep learning models like DeepBet which were trained explicitly on the macaque brain^30^. Other options focus more on image contrast quality and may include site-specific bias field correction and intensity normalization, both of which can improve surface reconstruction in precon_all. If the troubleshooting options above are insufficient, the most reliable path to a successful precon_all run is likely for the end user to optimize the quality of the input image directly.

### Troubleshooting common failure modes

After an initial run of precon_all, outputs are stored in a directory that follows the structure of FreeSurfer’s recon-all. This file structure eases the transition for users coming to non-human neuroimaging for the first time. As with FreeSurfer’s recon-all, the best way to inspect the cortical surface mesh output is visually, overlaying the white and pial surfaces on the brain-extracted image in Freeview. Users can also generate automated quality-control PDF reports using the *precon_qc.sh* script, which allow errors in brain extraction and pial surface placement to be quickly identified. Precon_all provides multiple fallbacks for each of these failure points (Table S1). Brain extraction failures are unique to the template-based workflow, since it relies on registration to transform a brain mask into subject space and extract the brain. If the FNIRT-based registration fails, users can instead try the ANTs-based option; if both fail, a manually edited brain mask can be supplied directly to precon_all using the -n flag, bypassing the native brain extraction step entirely.

Errors in surface reconstruction most often result from insufficient image contrast or intense inhomogeneity field insufficiently corrected by N4. This can cause the pial surface to expand outside the brain and can be corrected using *intensity_afni_unifize.sh*, which renormalizes image intensity and allows precon_2 or precon_3 to be rerun with the improved image, often resolving the issue (Fig. 3). Pial placement may also fail in that it does not sufficiently expand, typically affecting areas such as the temporal pole or a specific gyrus or sulcus missed during white matter segmentation (Fig. 2). As with FreeSurfer, this is corrected by manually editing the white matter segmentation mask generated by precon_2. Editing the mask by hand ensures the full extent of the white matter is captured, giving the pial surface an accurate starting point from which to expand in accordance with the anatomy present in the T1w image. Following manual editing, users can reconstruct the surface directly with the new white matter segmentation by rerunning precon_3.

### Building group-level surface templates

Precon_all can also be used to build group surface-based templates for spherical registration, extending the ability to capture group statistics of surface-based measures from human to preclinical neuroimaging (Fig. 5). To create a surface template space for a new species, we recommend that users start by creating an anatomical T1-weighted volumetric template, and that they then run precon_all on that image. Doing so ensures that the geometry of the surface and volumetric templates remain consistent, facilitating the projection of spatially normalized volumetric data to the average surface space. The process of average template creation begins by specifying the output of precon_all run on a single subject or volumetric template. A registration template based on the specified subjects’ sulcal curvature and folding is created, and a list of subjects is passed to precon_all’s *“pet_sounds.sh”* script which registers them to the sulcal template. The template is then updated, and this process is iterated an additional three times. Once completed, *pet_sounds.sh* creates an average surface by calling the associated *average_surface_maker.sh* script. As an example, we show the output of the average surface construction in the pig and macaque (Fig. 5), and note that as in human neuroimaging, these template spaces enable the spatial normalization of multimodal data to the common surface space.

### Surface registration

Precon_all does not perform surface registration out of the box, but includes a utility script (*register_surfaces.sh*) that allows users to register subjects to one another, or to a common surface template space, once precon_all has been run.

### Code Availability

All code is open source and freely available (https://github.com/neurabenn/precon_all) and can be installed by following the documentation, or run through a docker container (https://hub.docker.com/r/neurabenn/precon_all). Surfaces used in Fig. 4 including the pig, chimpanzee, and macaques are available upon request, while the remaining primate and carnivore surfaces can be found through the digital brain zoo ^7,17^.

## Acknowledgements

Ministry of Science, Innovation and Universities (PID2019-107332RB-I00 and PID2022-140176OB-I00), Instituto de Salud Carlos III (ISCIII; PI16/02110), and Comunidad de Madrid (S2017/BMD-3867 and P2022/BMD-7403 RENIM-CM). RAB was supported by a fellowship from the FP7-PEOPLE-2013-ITN. “Cardionext”. B.I is recipient of a European Research Council grant MATRIX (ERC-COG-2018-ID: 819775). ED received funding from SSNAP “Support for Sick and Newborn Infants and their Parents” Medical Research Fund (University of Oxford Excellence Fellowship). MB is a recipient of the L’ORÉAL Austria fellowship within the initiative “For Women in Science” and her research was funded in part by the Austrian Science Fund (FWF) 10.55776/J4828. DSM is supported by the European Research Council (ERC) under the European Union’s Horizon 2020 research and innovation programme (Grant agreement No. 866533). RBM is supported by the Biotechnology and Biological Sciences Research Council (BBSRC) UK [BB/N019814/1] and by the Medical Research Council (MRC) UK [MR/Y010698/1]. The Wellcome Centre for Integrative Neuroimaging is supported by core funding from the Wellcome Trust [203139/Z/16/Z]. The CNIC is supported by the Instituto de Salud Carlos III (ISCIII), the Ministerio de Ciencia, Innovación y Universidades and the Pro CNIC Foundation and is a Severo Ochoa Center of Excellence (grant CEX2020-001041-S funded by MICIN/AEI/10.13039/501100011033). For the purpose of Open Access, the author has applied a CC BY public copyright licence to any Author Accepted Manuscript version arising from this submission. We would also like to thank Andrew Rouse, for his help debugging the precon_all docker image.

## Author contributions

R.A.B. designed the processing pipeline, wrote the code, and wrote the manuscript. T.X. provided code for the surface generation steps generalized from precon_all and contributed to the manuscript. R.B.M. contributed data, proposed features central to precon_all, supervised the work, and contributed and aided in writing and revising the manuscript. M.B. provided the carnivoran surface data, contributed code to the repository, and contributed to the manuscript. L.R. provided data, tested the pipeline, and provided pipeline documentation. R.T. D.S.M, and J.P.M. contributed to writing and revision of the manuscript. P.M., C.G.A., and G.L.M. acquired data used in the study and contributed to the manuscript. E.P. supervised the work and contributed to the manuscript. B.I. contributed to the study design, supervised the project, and acquired funding.

