## Supplementary materials for "Precon_all: A species-agnostic automated pipeline for non-human cortical surface reconstruction"

**Supplementary Table S1. Software options and manual fallback points in precon_all**

| **Processing step** | **Template-based run** | **Single-subject / standard run** | **Manual or custom-module fallback** |
| --- | --- | --- | --- |
| Brain extraction & template registration | FSL FNIRT (legacy) or ANTs (-e); template tissue probability maps can be propagated to the individual image using the calculated warps | Brain extraction performed without template-based tissue priors | Supply an image that has already been brain extracted, using -n; extraction may come from another software package, a custom model (e.g., U-NET), or a manually defined mask |
| Tissue segmentation | FSL FAST or ANTs Atropos; template-derived tissue probability maps may optionally be used as priors | FSL FAST or ANTs Atropos, without template-derived priors | Supply a manually or externally generated binary white-matter segmentation as white_matter_hand_edit.nii.gz in the mri/ directory |
| Intensity normalization (fallback) | AFNI 3dUnifize | AFNI 3dUnifize | Images can be reprocessed using 3dUnifize and the pipeline rerun from the segmentation stage |

### **Supplementary Figures:**

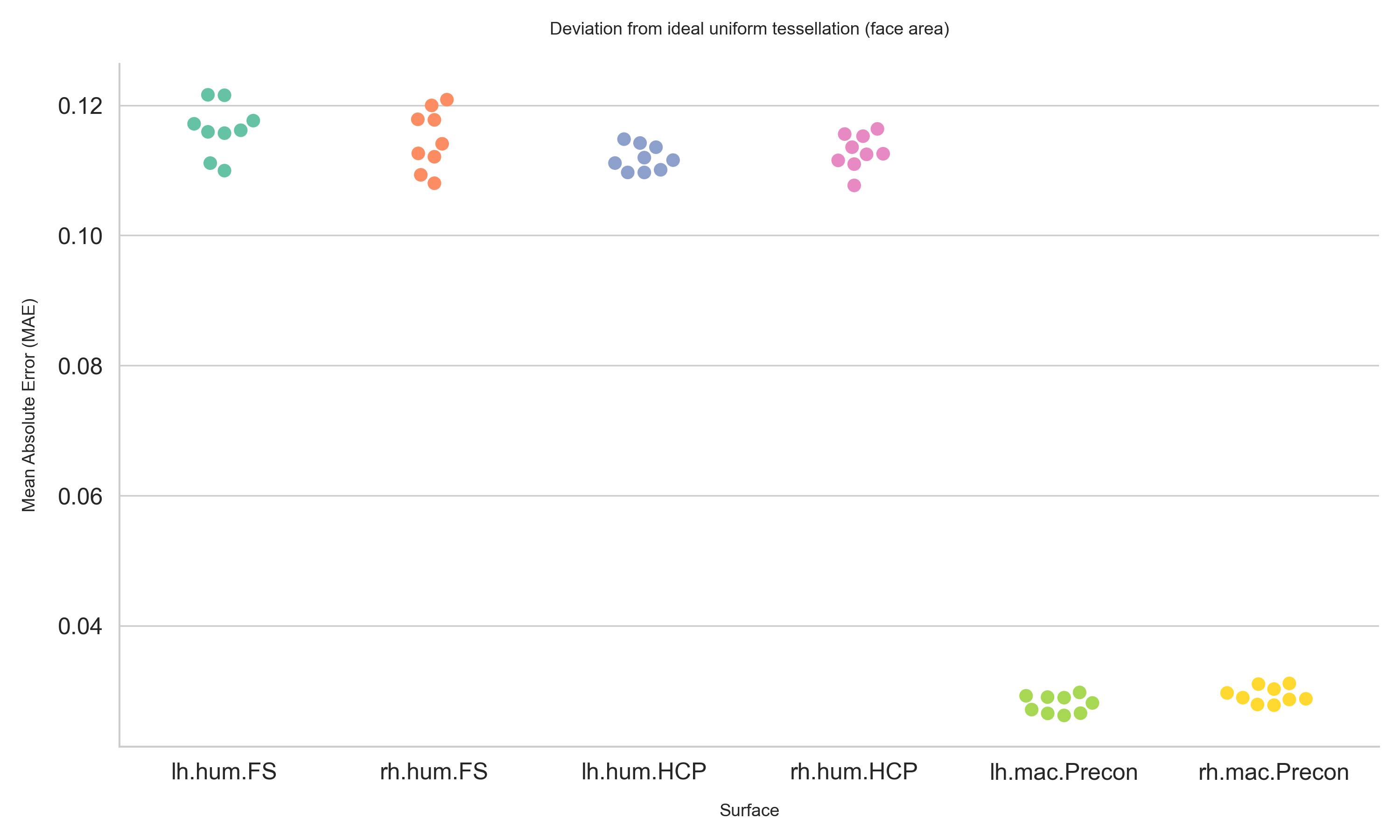

**Fig. S1 |** Comparison of Mean Absolute Error in Mesh Tessellations: This figure presents the mean absolute error between the ideal mesh tessellations and the actual results. Data include nine Human Connectome Project (HCP) subjects processed with either standard FreeSurfer or HCP pipelines, and nine macaques processed using Precon_all. Lower values indicate better performance, reflecting less error from an ideal uniform mesh. FreeSurfer meshes were generated on images downsampled to 1 mm isotropic resolution, while HCP meshes were processed at 0.7 mm isotropic. The macaque images were processed at 0.5 mm isotropic resolution.

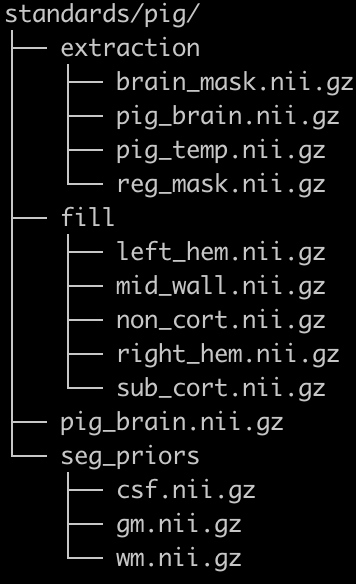

**Fig. S2 |** An example file structure used to automate precon_all for multiple subjects, where *“pig”* can be substituted for a user's species of choice.
